# IL-10-producing B cells are enriched in murine pericardial adipose tissues and ameliorate the outcome of acute myocardial infarction

**DOI:** 10.1101/520742

**Authors:** Lan Wu, Rajeev Dalal, Connie Cao, J. Luke Postoak, Qinkun Zhang, Zhizhang Wang, Hind Lal, Luc Van Kaer

## Abstract

Acute myocardial infarction (MI) provokes an inflammatory response in the heart that removes damaged tissues to facilitate repair. However, exaggerated and/or persistent inflammation compromises healing, which may be counteracted by regulatory immune mechanisms. A key regulatory factor in an inflammatory response is the anti-inflammatory cytokine IL-10, which can be produced by a number of immune cells including subsets of B lymphocytes. Here, we investigated IL-10-producing B cells in pericardial adipose tissues (PATs) and their role in the healing process following acute MI in mice. We found abundant IL-10-producing B cells in PATs under homeostatic conditions, with the majority of them bearing cell surface CD5 (CD5^+^ B cells). These cells were detected early in life, maintained a steady presence during adulthood, and resided in fat-associated lymphoid clusters (FALCs). The cytokine IL-33 was preferentially expressed in PATs under homeostatic conditions and contributed to enrichment of IL-10-producing CD5^+^ B cells in PATs. CD5^+^ B cells expanded in PATs following MI, and accumulated in the infarcted heart during the resolution of MI-induced inflammation. B cell-specific deletion of IL-10 worsened cardiac function after MI, exacerbated myocardial injury, and delayed resolution of inflammation. These findings reveal a significant contribution of IL-10-producing B cells to the anti-inflammatory mechanism that terminates MI-induced inflammation, and identify these cells as novel therapeutic targets to improve the outcome of MI.

**Significance Statement:** Myocardial infarction (MI) remains a leading cause of mortality and morbidity worldwide. Although it is now recognized that a balanced and timely terminated pro-inflammatory response following acute MI is essential in promoting tissue repair, the underlying regulatory mechanisms are poorly defined. In this report, we show that IL-10-producing B cells in mice 1) are enriched in pericardial adipose tissues (PATs) and influenced by cytokine IL-33 under homeostatic conditions; 2) expand in PATs following MI and accumulate in the infarcted heart during the resolution of MI-induced inflammation; and 3) facilitate resolution of inflammation and reduce myocardial injury to preserve cardiac function after MI. These findings identify IL-10-producing B cells as novel therapeutic targets to improve the outcome of MI.

## Introduction

Despite advances in prevention and treatment, acute myocardial infarction (MI) remains a leading cause of mortality and morbidity worldwide (1, 2), highlighting the need for new therapeutic strategies. Acute MI provokes a sterile inflammatory response that centers in the heart and involves primarily the innate branch and to a lesser extent the adaptive branch of the immune system (3–5). The early “inflammatory phase” is characterized by recruitment of pro-inflammatory cells and release of pro-inflammatory mediators. Although these events are critical for instigating tissue repair by removing damaged tissues, exaggerated and/or persistent pro-inflammatory reactions exacerbate myocardial injury and contribute to short- and long-term complications (6–8). As such, a balanced inflammatory phase and its timely termination move the healing process to the subsequent “proliferative phase” followed by a “maturation phase” when the tissue environment in infarcted hearts transitions to an anti-inflammatory state to promote a reparative process. While recent studies have begun to investigate the underlying mechanisms (9–12), much remains to be learned regarding the cells and molecules that regulate the progression of MI-induced inflammation. In this context, several reports have identified a beneficial role of the anti-inflammatory cytokine IL-10 in the outcome of acute MI (13–17). However, the cellular sources and downstream targets of this cytokine that collectively provide cardioprotection remain to be defined.

Mature B cells can be classified into B-1 and B-2 cells based on a combination of surface phenotype and functional properties, and each class of B cells is comprised of distinct subsets (18–20). B-2 cells are also referred to as conventional B cells and make up the majority of B cells in the primary and secondary lymphoid organs. B-1 cells belong to a group of so-called innate-like B cells that respond well to innate signals such as toll-like receptor ligands and other conserved pathogen-derived molecular patterns (21). Surface expression of CD5 further distinguishes B-1a (CD5^+^) from B-1b (CD5^-^) cells. In addition to their well-known function to produce antibodies, B cells fulfill a number of antibody-independent functions. Among the latter B cell properties is the release of cytokines that in turn regulate inflammation (22, 23). Subsets of B cells that release either stimulatory or inhibitory cytokines have been identified (22). Published studies have implicated a pathogenic role of B cell subsets producing pro-inflammatory cytokines in acute MI (24, 25). While still under debate, the term regulatory B cells has been employed in recent years to describe B cell subsets that modulate inflammation through production of inhibitory cytokines, with IL-10-producing B cells as the best studied subset (26–28). Judging by their surface phenotype and response to stimuli, which can resemble either B-1 or B-2 cells, it is likely that IL-10-producing B cells differentiate along the developmental pathway for either the B-1 or the B-2 cell lineages (29, 30). CD5^+^ B-1a cells are the main IL-10-producing B cells in the peritoneal cavity (PerC) where subsets of B-1 cells preferentially reside (31, 32), whereas IL-10-producing CD5^+^ B10 cells phenotypically resemble B-2 cells (33). We and other investigators recently reported that murine abdominal visceral adipose tissues (VATs) represent another site where B-1 cells preferentially reside. These adipose depots contain a significant pool of IL-10-producing B cells that protect against obesity-induced inflammation, with the majority of them bearing a surface phenotype of B-1a cells (34, 35).

VATs are also present in the thoracic cavity. The adipose depots associated with the heart include epicardial adipose tissues (EATs) located between the myocardium and visceral pericardium (epicardium) or around coronary arteries, and pericardial adipose tissues (PATs) located between the epicardium and parietal pericardium (36). The presence of significant quantities of both EATs and PATs in humans and large animals has been long recognized (37–39). In mice, EATs are scant but not absent, whereas PATs are readily discernible (39). It has been known that PATs house aggregates of leukocytes (40–42). More recently, these anatomical structures have been referred to as fat-associated lymphoid clusters (FALCs) (43–45) that belong to a group of physiological and inflammation-induced tertiary lymphoid tissues (46, 47). Detailed analyses of FALCs in murine mesenterium revealed a predominance of B cells, with a large proportion of B-1 cells (44). In response to peritoneal inflammatory stimuli, FALCs of abdominal VATs drastically expand to support B cell proliferation with a preferential expansion of B-1 cells (44). Previous studies showed a high density of FALCs in murine PATs under homeostatic conditions (44, 45). A recent report provided evidence for expansion of these lymphoid tissues following acute MI (25). However, the composition of B cell subsets in FALCs of PATs and their response to MI-induced inflammation have not been explored. Moreover, IL-10-producing B cells in FALCs of PATs under homeostatic conditions and their role in acute MI remains unclear. We report here that IL-10-producing B cells are abundant in murine FALCs of PATs under homeostatic conditions, with the majority of them bearing the surface phenotype of CD5^+^ B-1a cells. Our studies further show that these cells play a cardioprotective role after acute MI.

## Results

### PATs house abundant IL-10-producing CD5^+^ B cells under homeostatic conditions

We previously reported that IL-10-producing B cells were enriched in the B cell compartment of murine perigonadal VATs. The majority of them displayed a surface phenotype of CD5^+^ B-1a cells (34). To assess whether this is reflected in adipose tissues throughout the body, we analyzed a variety of adipose depots, including subcutaneous brown adipose tissues (BATs), subcutaneous white adipose tissues (SATs), and VATs in the two body cavities. We harvested interscapular BATs, inguinal SATs, abdominal (perigonadal, retroperitoneal, and mesenteric) VATs, and thoracic VATs (PATs and periaortic VATs, Figure 1a) from adult wild-type C57BL/6J (WT B6) mice. We first analyzed B cell subsets in purified stromal vascular fraction cells (SVFs) by flow cytometry, and compared them to cells harvested from PerC (Supplemental Figure 1a). The results showed that, in comparison to BATs and SATs, all depots of VATs had a higher prevalence of B-1 cells that were split roughly 1:1 between B-1a and B-1b cells (Supplemental Table 1). Because CD5^+^ B cells have a higher propensity of producing IL-10, and because we previously showed that CD5^+^ B cells made up the majority of IL-10-producing B cells in perigonadal VATs (34), we next quantified CD5^+^ B cells (Supplemental Figure 1b). The results showed that the prevalence of CD5^+^ B cells was significantly higher in depots of VATs than those in BATs or SATs (Figure 1b). The highest frequency (Figure 1b), together with a large B cell compartment (Supplemental Table 1), translated into a strikingly higher density of CD5^+^ B cells in PATs as compared with perigonadal VATs (Figure 1c).

**Figure 1.**
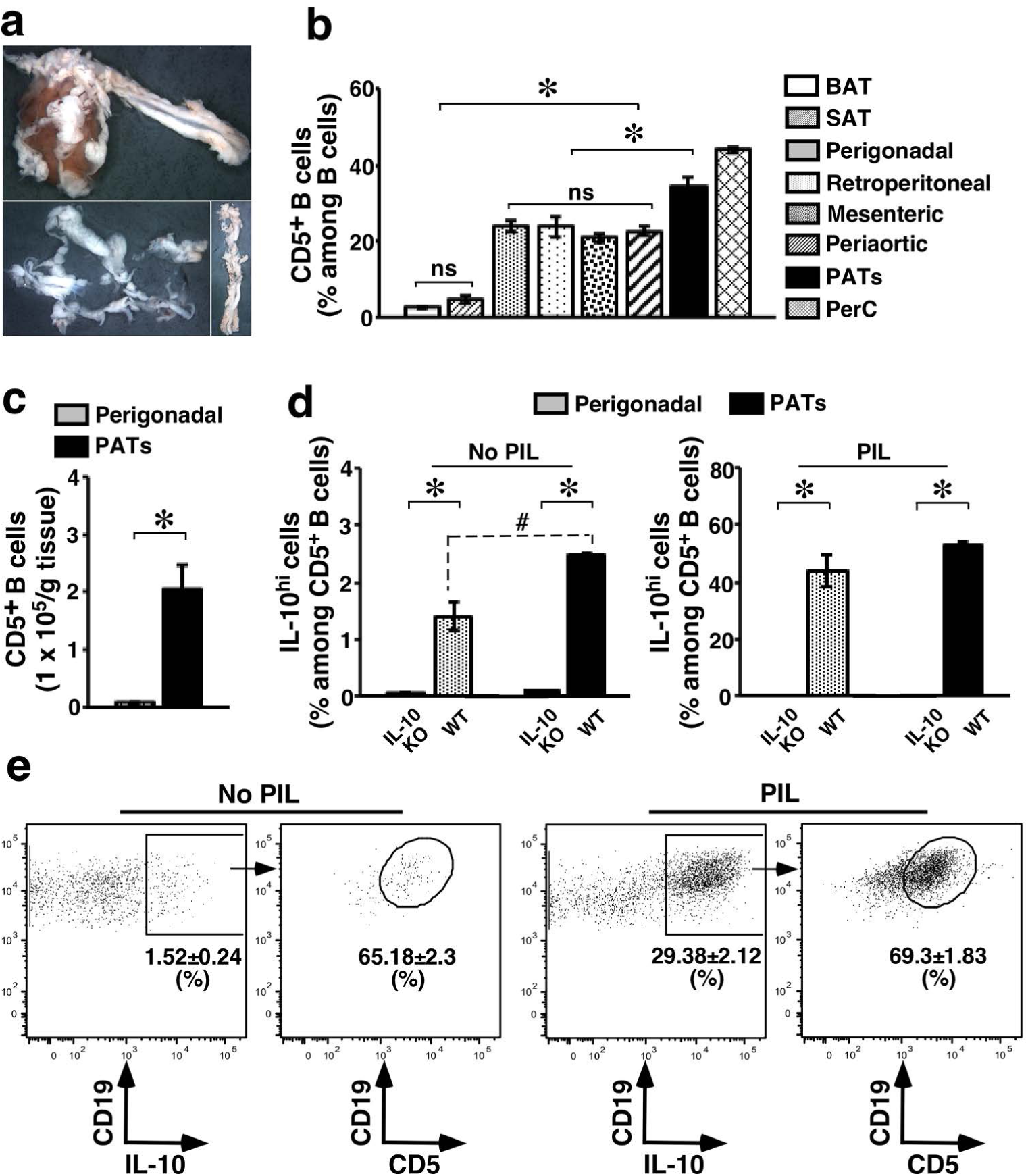
IL-10-producing CD5^+^ B cells in PATs. **a**. The heart and thoracic aorta with surrounding VATs from an adult WT B6 mouse were harvested and photographed (upper panel). PATs together with pericardium (lower left) or periaortic VATs (lower right) were separated from heart or thoracic aorta, and then photographed (magnification: 0.5x). **b**. SVFs from the indicated adipose depots were examined by flow cytometry for the prevalence of CD5^+^ B cells (summary of 2 independent experiments, n=8). **c**. The tissue density of CD5^+^ B cells in the indicated VAT depots is shown (summary of 2 independent experiments, n=8). **d**. SVFs from the indicated VAT depots of mice with the indicated genotypes were cultured for 5 hours in the presence of monensin, with or without the mitogen mixture PIL (PMA, ionomycin, and LPS). Cells were then examined by flow cytometry for IL-10^hi^ CD5^+^ B cells. A summary of 2 independent experiments is shown (n=6). **e**. SVFs from PATs of WT B6 mice were treated as described in **d** and examined by flow cytometry. IL-10^hi^ B cells were further analyzed for surface CD5 expression. Representative flow cytometry plots from 2 independent experiments are shown (n=6). ^#^: p<0.05; *: p<0.01; and ns: not statistically different for all panels.

We next examined IL-10 production, and compared cells harvested from PATs with those from perigonadal VATs. Among B cells freshly isolated from PATs of WT B6 mice, we detected very little IL-10 when comparing mean fluorescence intensity (MFI), and the prevalence of cells expressing high levels of IL-10 (IL-10^hi^) was negligible (Supplemental Figure 2a). We used cells from IL-10 knockout mice (IL-10 KO) as a control to further ascertain intracellular IL-10. In CD5^+^ B cells isolated from the two VAT depots of WT B6 mice, IL-10 accumulated to a detectable level after a short-term *ex vivo* culture in the presence of the protein transport inhibitor monensin. This was reflected by increases in the frequency of IL-10^hi^ cells (Figure 1d, left panel). A short-term *ex vivo* challenge with a mixture of B cell mitogens, PIL (phorbol myristate acetate [PMA], ionomycin, and lipopolysaccharide [LPS]), further promoted IL-10 production (Figure 1d, right panel). Analysis of the MFI of IL-10 staining confirmed these observations (Supplemental Figure 2b). These results indicate that CD5^+^ B cells in PATs can function as IL-10-producing cells, in a manner similar to but more potent than their counterparts in perigonadal VATs (Figure 1d and Supplemental Figure 2b). These cells continuously produce low levels of IL-10 under unchallenged conditions and further enhance IL-10 production upon stimulation. Similar to our previous observations in perigonadal VATs, CD5^+^ B cells constituted the majority of IL-10-producing B cells in PATs and this was true under both unchallenged and PIL-challenged conditions (Figure 1e). Most of these cells displayed a surface phenotype resembling B-1a cells rather than B-2 cells (Supplemental Figure 3). Therefore, the majority of IL-10-producing CD5^+^ B cells in adipose tissues likely belong to the B-1a cell lineage. Collectively, the above findings demonstrate the presence of abundant IL-10-producing B cells in PATs under homeostatic conditions with the majority of them bearing surface CD5.

**Figure 2.**
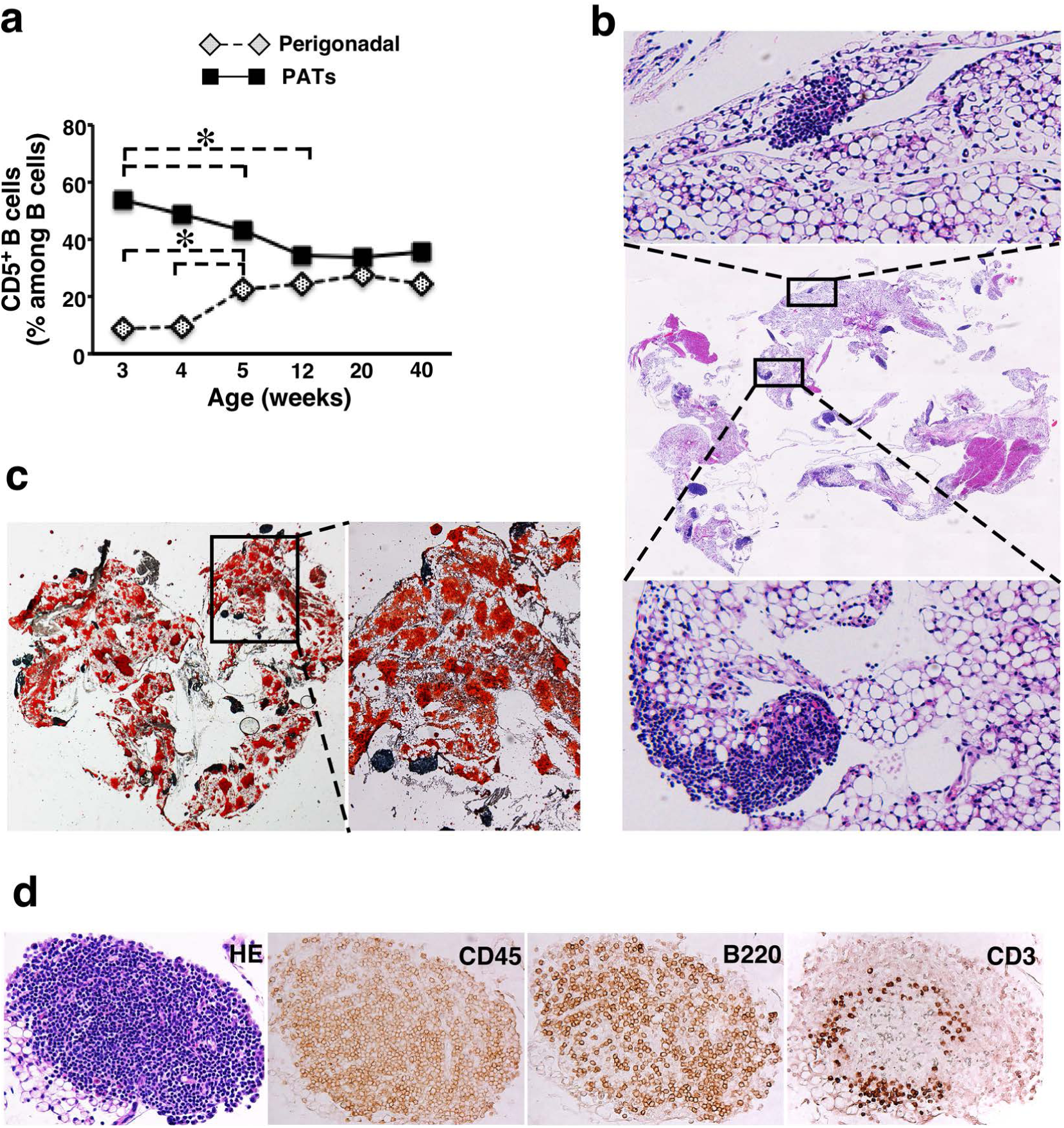
Population dynamics and residence of CD5^+^ B cells in PATs. **a.** Prevalence of CD5^+^ B cells in the indicated locations was examined in WT B6 mice at different ages. A summary of 2 independent experiments (n=8 for each age group) is shown (*: p<0.01). **b**. Paraffin-embedded sections of PATs together with pericardium were stained with H&E and photographed (magnification: 0.5x). Two areas with densely packed cells are enlarged to show clusters (magnification: 4x). Representative graphs from 3 independent experiments are shown (n=6). **c**. Frozen sections of PATs together with pericardium were stained with Oil Red O and photographed (magnification: 0.5x). One area with densely packed cells is enlarged to show clusters (magnification: 4x). Representative graphs from one experiment are shown (n=3). **d**. Serial paraffin-embedded sections were either H&E-stained or immunostained for the indicated markers and photographed. Images of a representative cluster are shown (magnification: 20x).

**Figure 3.**
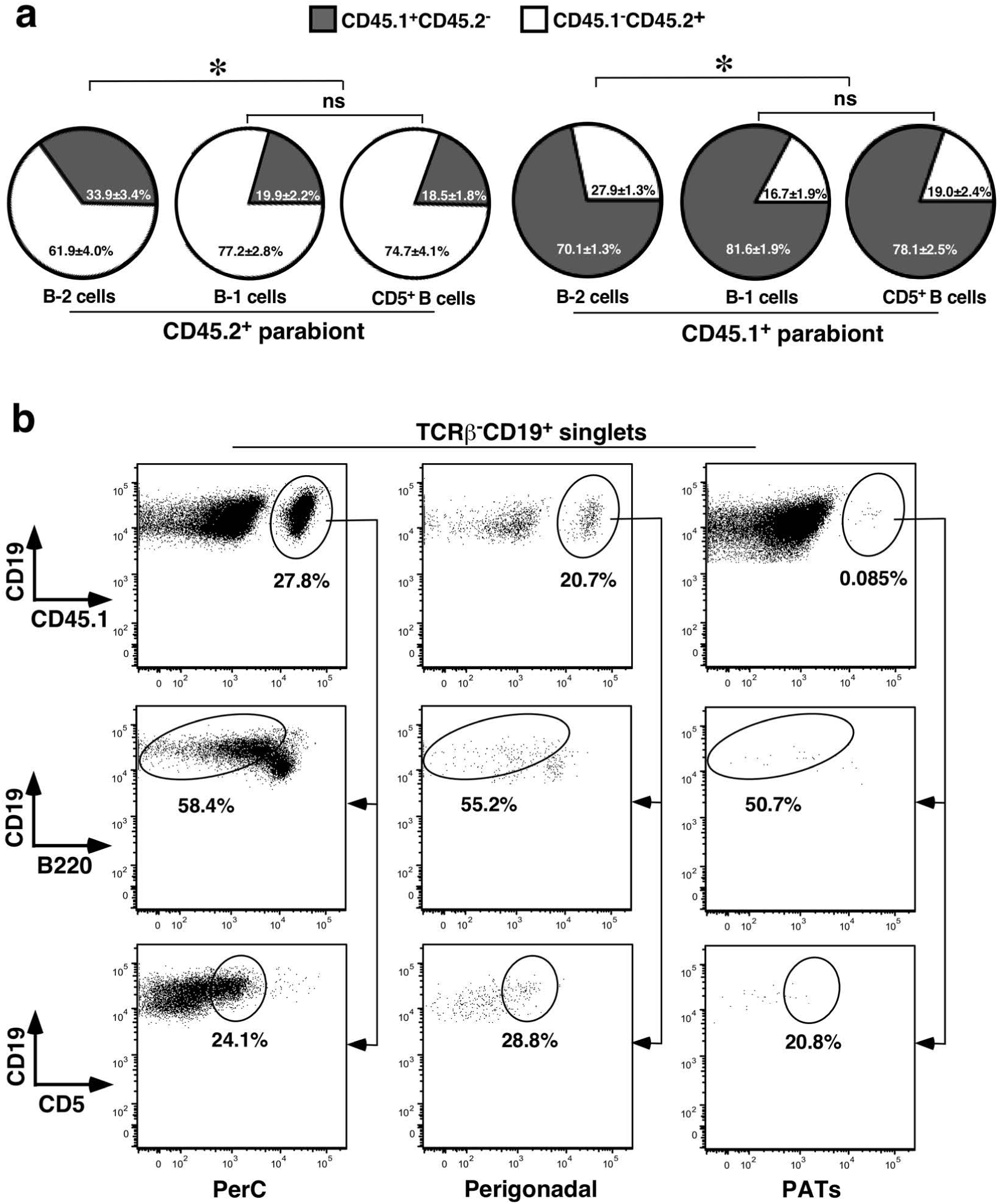
Maintenance of CD5^+^ B cells in PATs during adulthood. **a**. Congenic CD45.1^+^ and CD45.2^+^ mice were joined by parabiosis and examined one month later. Subsets of B cells in PATs of each parabiont were examined by flow cytometry. Summary of 3 independent experiments with 11 pairs of mice is shown (*: p<0.01; and ns: not statistically different). **b**. Cells harvested from PerC of CD45.1^+^ mice were transferred into PerC of CD45.2^+^ mice. The indicated tissues were examined 1 week later for the prevalence of donor cells among different B cell subsets. Representative flow cytometry plots from 2 separate experiments are shown (n=6).

### CD5^+^ B cells seed PATs early in life, maintain a steady presence during adulthood, and reside in FALCs

To investigate the population dynamics of CD5^+^ B cells in PATs, we analyzed WT B6 mice at different ages. In PATs, CD5^+^ B cells were detected at the highest frequency when mice were 3 weeks of age. The prevalence of these cells gradually decreased as mice matured to reach steady-state levels (Figure 2a). The age-dependent population dynamics of CD5^+^ B cells appeared depot-specific. Seeding of CD5^+^ B cells in perigonadal VATs lagged behind PATs. The prevalence of these cells in perigonadal VATs increased as mice matured to reach steady-state levels during adulthood (Figure 2a). The prevalence of CD5^+^ B cells in thoracic periaortic VATs followed the pattern observed for perigonadal VATs whereas the frequency of these cells in PerC resembled the one found in PATs (Supplemental Fig. 4).

**Figure 4.**
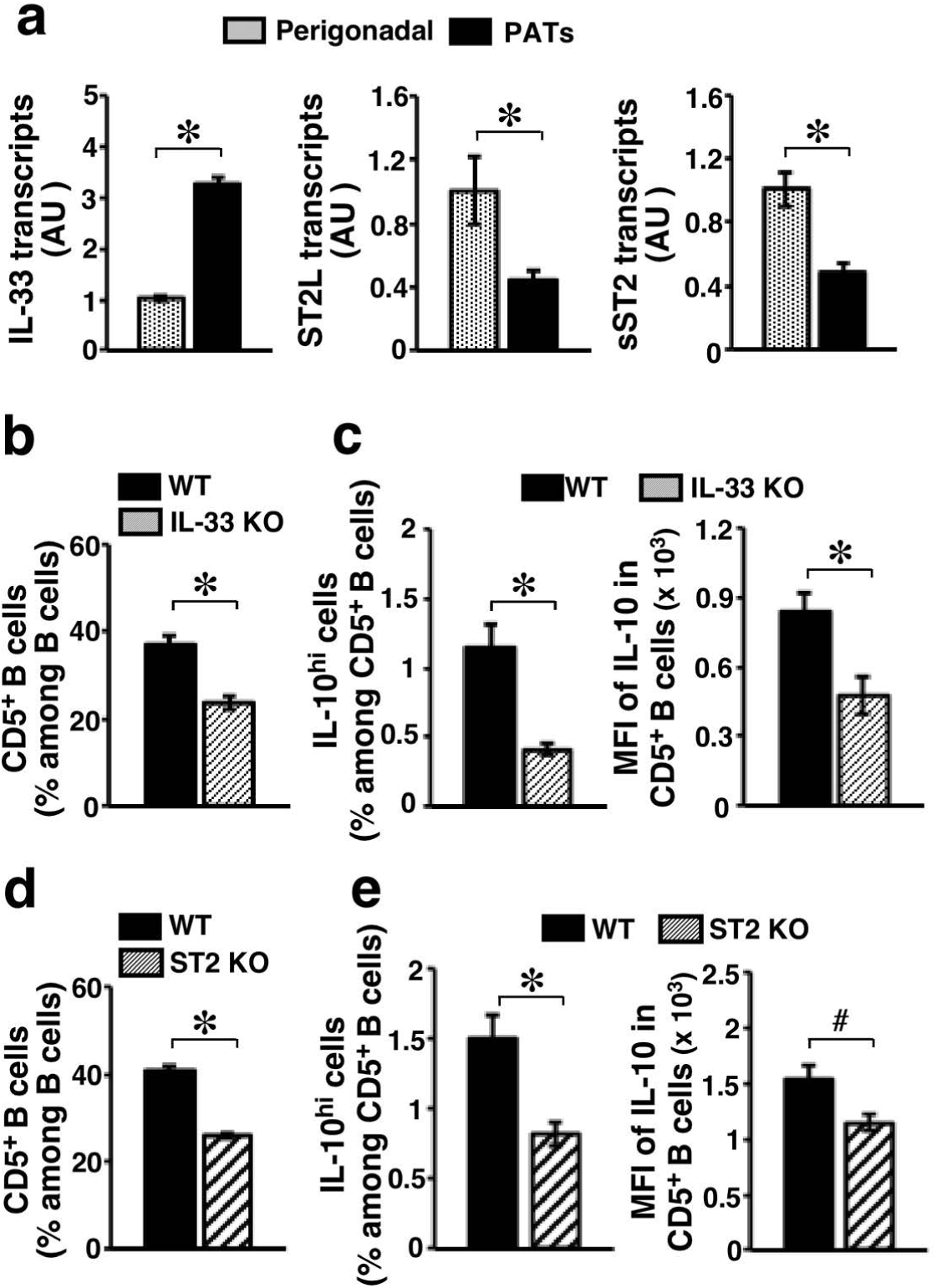
Influence of IL-33 signaling on CD5^+^ B cells in PATs. Adult mice of the indicated genotypes were sacrificed to harvest the tissues. **a**. Transcripts of the indicated genes were examined in the indicated VAT depots of WT B6 mice by real-time PCR. Summary of 2 independent experiments is shown (n=8; AU: arbitrary unit). **b, d**. SVFs from PATs of mice with the indicated genotypes were examined by flow cytometry for the prevalence of CD5^+^ B cells. **c, e**. SVFs from PATs of mice with the indicated genotypes were *ex vivo* cultured for 5 hours in the presence of monensin, and examined by flow cytometry for IL-10^hi^ CD5^+^ B cells (left panels) and the intensity of IL-10 staining in CD5^+^ B cells (right panels). Summary of 2 independent experiments is shown for **b**-**e** (n=9-12 for each group). ^#^: p<0.05; and *: p<0.01 for all panels.

We next examined the tissue residence of CD5^+^ B cells in PATs. We prepared whole-mount PATs together with pericardium (see Figure 1a) and stained the tissues with hematoxylin and eosin (H&E). As shown in Supplemental Figure 5a, dense clusters of cells in different sizes and shapes were readily detectable. On H&E-or oil red O-stained tissue sections, these clusters were embedded in or associated with adipose tissues (Figure 2, b and c). Similar to observations in humans (48), murine PATs were comprised of adipocytes with either unilocular (white adipocytes) or multilocular (brown adipocytes) morphology (Supplemental Figure 5b). The non-adipocyte clusters were always located in or associated with white adipose tissues (Figure 2b). The vast majority of cells inside clusters stained positive for CD45, indicating a hematopoietic origin, with B cells significantly outnumbering T cells and populating the entire clusters (Figure 2d). Dual immunofluorescence staining for CD45 and a B cell marker confirmed these findings (Supplemental Figure 5c). These results demonstrate that the densely packed clusters of cells in PATs are indeed FALCs. Of note, formation of B cell-dominant FALCs in PATs does not appear to require IL-10, as evidenced by their presence in IL-10 KO mice (Supplemental Figure 6).

**Figure 5.**
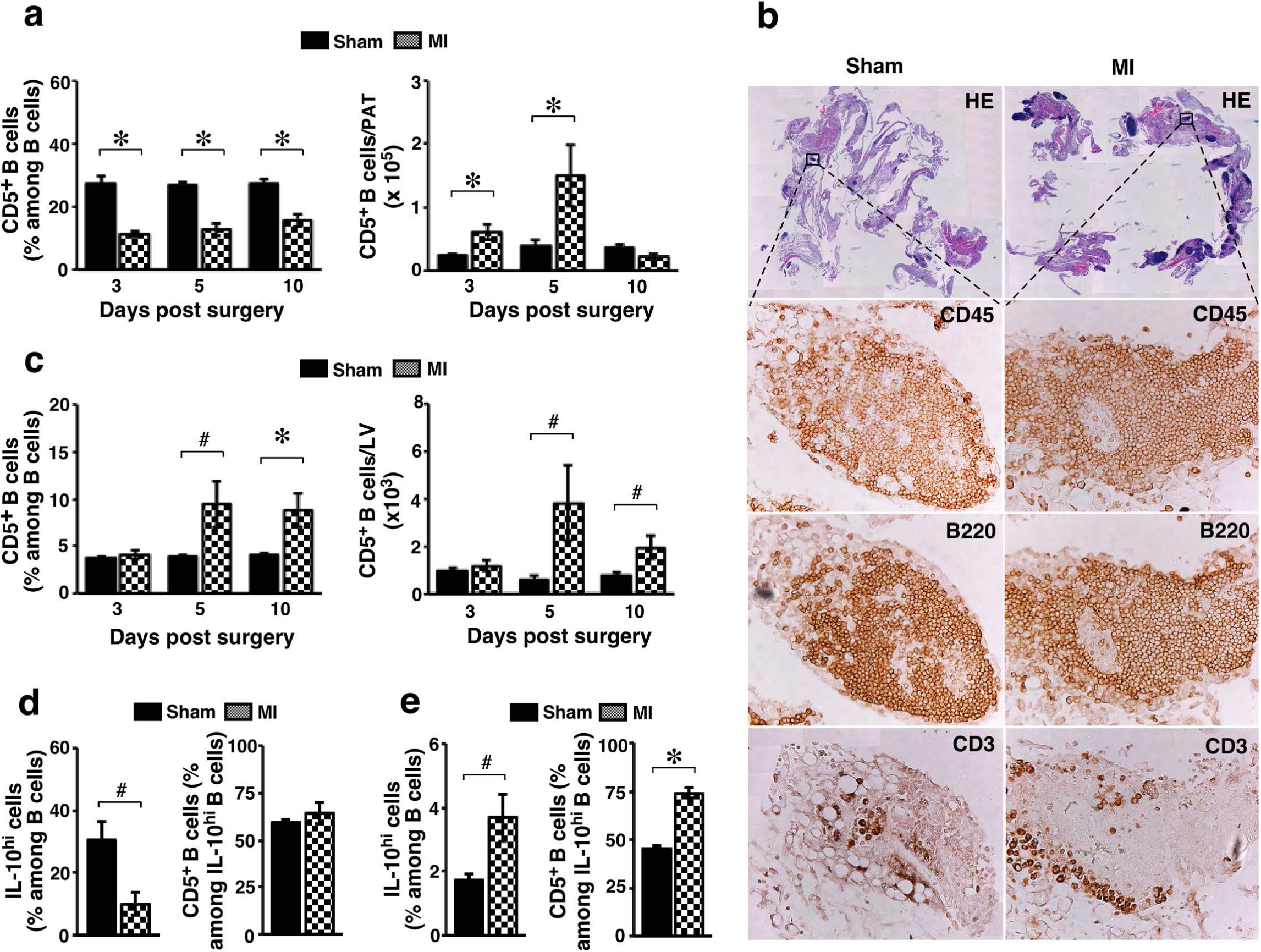
MI-induced expansion of CD5^+^ B cells in PATs following MI and their accumulation in the infarcted hearts. **a.** WT B6 mice underwent sham or MI surgery and were analyzed at the indicated time post-surgery. SVFs from PATs were examined for CD5^+^ B cells (summary of 2-3 independent experiments, n=8-10 for each group at each time point). **b**. Sham- or MI-operated WT B6 mice were examined 5 days post-surgery. Serial paraffin-embedded sections of PATs together with pericardium were stained and photographed (magnification: 0.5x). FALCs were immunostained for the indicated markers and photographed (magnification: 20x). Representative graphs from 2 independent experiments are shown (n=4). **c**. Mice were treated as described in **a**. Leukocytes from LVs were examined for CD5^+^ B cells (summary of 2-3 independent experiments, n=8-10 for each group at each time point). **d**, **e**. WT B6 mice underwent sham or MI operation and were analyzed 8 days post-surgery. SVFs from PATs (**d**) or leukocytes from LVs (**e**) were *ex vivo* challenged. Cells were then examined by flow cytometry for intracellular IL-10 (summary of 2 independent experiments, n=4-6 per group). ^#^: p<0.05; and *: p<0.01.

**Figure 6.**
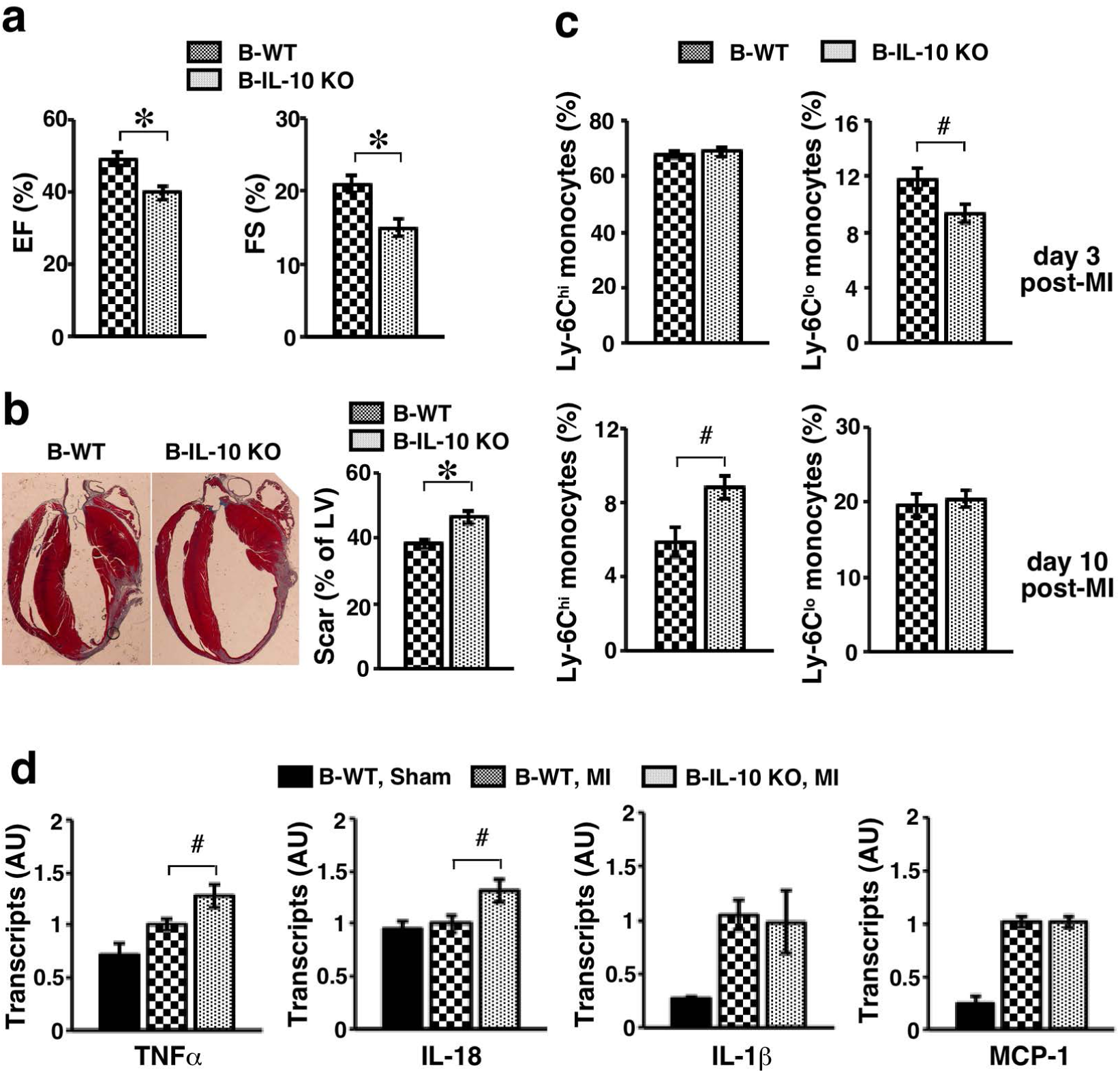
Impact of B cell-specific deletion of IL-10 on the outcome of acute MI. B-IL-10 KO and B-WT mice underwent MI surgery. **a**. Mice were examined for LV function by echocardiography 3 weeks post-MI. EF (left) and FS (right) are shown (n=13-17 in each group). **b**. Mice were sacrificed 3 weeks after surgery. Representative images of Masson’s trichrome stain are shown in the left panel (magnification: 0.5x). Scar areas were measured and are presented as % of LV circumference in the right panel (n=17 in each group). **c**. Mice were sacrificed at the indicated time post-MI to purify leukocytes from LVs. Subsets of monocytes were analyzed by flow cytometry. Summary of 2 independent experiments (n=8 for each group at each time point) are shown. **d**. Transcripts of the indicated genes were examined in LVs harvested 2 weeks post-MI by real-time PCR. Summary of 2 independent experiments is shown (n=7-11). ^#^: p<0.05; and *: p<0.01.

Previous studies on the subsets of B cells in PerC have revealed that B-1 cells are largely generated during fetal and neonatal development. The specified progenitors produce less B-1 cells, in particular B-1a cells, during adulthood in comparison to the B-2 lineage (49–51). We therefore examined whether this was the case for B cell subsets in PATs. We performed parabiosis using congenic B6 mice expressing either CD45.1 or CD45.2 congenic markers to track cells, and analyzed subsets of B cells in conjoined pairs. The results showed that, after one month of parabiosis, a sizable fraction of CD5^+^ B cells in PATs was derived from their parabiotic partner. This fraction was, however, significantly smaller than the B-2 cells (Figure 3a). Similar results were obtained when B-1 cells were analyzed (Figure 3a). These observations held true for perigonadal VATs (Supplemental Figure 7). As such, the maintenance of B cell subsets in VATs during adulthood appears consistent with that in PerC. Because parabiosis creates a shared circulation between joined partners, one potential source for the new recruitment is re-circulated CD5^+^ B cells as opposed to those supplied by progenitors. To distinguish between these possibilities, we collected B cells from PerC of CD45.1^+^ mice and adoptively transferred them into PerC of CD45.2^+^ mice. While transferred CD5^+^ B cells migrated to perigonadal VATs, few of these cells had migrated to PATs 7 days post-transfer (Figure 3b). Adoptive transfer of B cells harvested from PATs into PerC of recipient animals yielded similar results. Therefore, the CD5^+^ B cell compartments in the abdominal and thoracic cavities appear well segregated, lending support for the scenario that progenitors supply CD5^+^ B cells to PATs during adulthood. Together, the above findings demonstrate that abundant CD5^+^ B cells reside in FALCs of PATs under homeostatic conditions throughout life.

### Preferential expression of IL-33 in PATs contributes to the enrichment of and IL-10 production by CD5^+^ B cells under homeostatic conditions

To investigate the mechanisms involved in the enrichment of and IL-10 production by CD5^+^ B cells in PATs, we examined the influence of the cardioprotective cytokine IL-33 (52, 53), which has been shown to influence B-1 cells (45, 54). Binding of IL-33 with the membrane-bound receptor ST2 (ST2L) transduces signals in target cells, whereas sequestration of this cytokine by secreted soluble ST2 (sST2) blocks its effects (52). In WT B6 mice, the expression of IL-33 was significantly higher and expression of ST2 isoforms was significantly lower in PATs compared to perigonadal VATs (Figure 4a). To ascertain whether the IL-33/ST2L signaling axis indeed influenced CD5^+^ B in PATs, we utilized mouse models of IL-33- or ST2-deficiency (55, 56). IL-33-deficiency (IL-33 KO) in mice caused disturbances in the subset distribution of B cells in PATs under homeostatic conditions, with significant decreases in B-1a cells (Supplemental Figure 8a). Consequently, the prevalence of CD5^+^ B cells was significantly reduced (Figure 4b). The results for ST2-deficient (ST2 KO) mice were consistent with those for IL-33 KO mice (Figure 4d, and Supplemental Figure 8b). IL-10 production in the remaining CD5^+^ B cells was significantly lower in both IL-33- and ST2-deficient mice (Figure 4, c and e). These results indicate that locally produced IL-33 acts through ST2 to enhance CD5^+^ B cells in PATs under homeostatic conditions.

### CD5^+^ B cells expand in PATs following acute MI and accumulate in infarcted hearts during the resolution phase of MI-induced inflammation

To assess the response of CD5^+^ B cells during acute MI, we employed a mouse model of acute MI by surgically ligating the left anterior descending coronary artery (LAD) (57). Similar to human pathology, such manipulation provokes an inflammatory response in the heart (3–5). We performed a sham or MI procedure on WT B6 mice, and analyzed PATs and left ventricles (LVs) at 3, 5, and 10 days post-surgery to sample the transition from a pro-inflammatory to an anti-inflammatory tissue environment (58). In PATs, acute MI caused a decrease in the percentage of CD5^+^ B cells among B cells during the entire time period examined (left panel in Figure 5a). This may reflect expansion of B-2 cells that outpaces expansion of B-1 cells in response to acute MI (49–51). Nevertheless, the total number of CD5^+^ B cells per PAT was significantly increased at 3 and 5 days post-MI (right panel in Figure 5a), indicating an MI-induced expansion of CD5^+^ B cells that peaks during the transition to the resolution phase of inflammation. Histological analyses of PATs, both whole-mount (Supplemental Figure 9) and tissue sections (Figure 5b), confirmed the above findings by demonstrating an expansion of B cell-dominant FALCs.

Compared to PATs, the normal mouse heart contained a much smaller B cell compartment (Supplemental Table 1). The prevalence of B-1 cells, in particular B-1a cells, was significantly lower (Supplemental Figure 10a and Supplemental Table 1). CD5^+^ B cells only represented a minor subset of B cells in normal hearts (Supplemental Figure 10b). Following acute MI, CD5^+^ B cells accumulated in the infarcted hearts after the initial inflammatory phase. This was reflected by increases in their prevalence and total cell numbers (Figure 5c). The response of CD5^+^ B cells to acute MI was injury site-specific. Parallel analyses of perigonadal VATs or PerC did not recapitulate the dynamics observed in PATs and LVs.

Lastly, we examined IL-10 production. Similar to PATs under homeostatic conditions, the levels of IL-10 in freshly purified B cells were very low in both PATs and LVs of sham- or MI-operated mice (Supplemental Figure 11). However, the response of B cells to additional challenge displayed differential profiles for PATs and LVs. During the resolution phase of MI-induced inflammation, IL-10-producing B cells were significantly reduced in PATs (left panel in Figure 5d), but significantly increased in LVs of MI-operated mice (left panel in Figure 5e). The increases in ventricular IL-10-producing B cells were largely accounted for by increases in CD5^+^ B cells (right panel in Figure 5e). Together, our findings indicate that IL-10-producing CD5^+^ B cells expand in PATs after MI, and accumulate in infarcted hearts as the MI-induced response makes its transition to create an anti-inflammatory tissue environment.

### B cell-specific deficiency of IL-10 worsens cardiac function after MI, exacerbates myocardial injury and delays resolution of inflammation

We next investigated whether IL-10-producing B cells afford cardioprotection during acute MI. Because our studies showed that B-1a cells made up the majority but not the entire pool of adipose IL-10-producing B cells, we employed a loss-of-function model that deleted IL-10 in the B cell compartment. A mixed bone marrow chimera approach has been used for this purpose (59–62). We generated B6 mice containing IL-10-deficient B cells (B-IL-10 KO) and control mice containing IL-10-sufficient B cells (B-WT). After reconstitution of the transplanted bone marrow, we first confirmed the efficiency of this approach to delete IL-10 in B cells of PATs (Supplemental Figure 12) as well as of other VATs and PerC. We then performed MI surgery on mice from both groups. We employed echocardiography to examine the function of LVs 3 weeks post-MI (63). The results showed that lack of IL-10 production by B cells worsened LV function. This was reflected by a lower ejection fraction (EF, left panel in Figure 6a) and a lower fractional shortening (FS, right panel in Figure 6a), which were calculated from the parameters taken from M-mode images (Supplemental Figure 13). Masson’s trichrome staining of collagen showed larger scars in the hearts of B-IL-10 KO mice compared to B-WT mice, indicating exacerbated myocardial injury in the former animals (Figure 6b).

Acute MI provokes a sequential accumulation of two subsets of monocytes in infarcted hearts (9, 64). Pro-inflammatory Ly-6C^hi^ monocytes dominate the early phase to promote removal of damaged tissues, whereas accumulation of anti-inflammatory Ly-6C^lo^ monocytes is associated with transition to an anti-inflammatory tissue environment at the later phases. We therefore analyzed monocyte subsets in the infarcted LVs of B-WT and B-IL-10 KO mice. We employed a flow cytometry strategy (64) that was used in our previous studies on VATs (34). This analysis separates monocytes from other myeloid cells and further specifies the 2 subsets of monocytes. The results showed that influx of Ly-6C^hi^ monocytes (lower right quadrants in Supplemental Figure 14) was comparable between the two groups at 3 days post-MI (Figure 6c, upper left panel). This suggests that lack of IL-10 production by B cells does not impair the acute MI-induced pro-inflammatory response. However, the clearance of these cells was significantly delayed in B-IL-10 KO mice, as reflected by a higher prevalence of Ly-6C^hi^ monocytes 10 days post-MI (lower left panel in Figure 6c). Interestingly, the prevalence of Ly-6C^lo^ monocytes (lower left quadrants in Supplemental Figure 14) was significantly lower in B-IL-10 KO mice 3 days post-MI (upper right panel in Figure 6c), indicating a slower transition to the anti-inflammatory state. These findings implicate a regulatory role of IL-10-producing B cells in the MI-induced monocyte response. Finally, the overall tissue environment in the infarcted LVs remained more pro-inflammatory in B-IL-10 KO mice during the resolution of MI-induced inflammation, as reflected by higher levels of the pro-inflammatory cytokines TNF-α and IL-18 (Figure 6d). Collectively, these findings reveal that the presence of IL-10-producing B cells ameliorates the outcome of acute MI.

## Discussion

We report here that murine PATs house abundant IL-10-producing CD5^+^ B cells under homeostatic conditions. These cells continuously produce low levels of IL-10 under unchallenged conditions and further enhance production of this cytokine upon stimulation. In response to an acute MI, these cells significantly expand in PATs. They accumulate in the infarcted hearts during the resolution of MI-induced inflammation. The presence of IL-10-producing B cells improves the outcome of MI by preserving cardiac function and minimizing myocardial injury. The role of IL-10 in acute MI has drawn research attention in recent years. Despite some inconsistencies in the published findings (65), the accumulated evidence supports a beneficial role of this cytokine in improving the outcome of acute MI (13–17). Our studies now specify subsets of B cells as a cellular source for IL-10 that protects against MI-induced cardiac damage.

Our studies show that CD5^+^ B cells populate PATs early in life and are maintained there at a steady presence under homeostatic conditions. Based on the surface phenotype and a rapid response to MI-induced sterile inflammation, the majority of CD5^+^ B cells likely differentiate along the developmental pathway for the B-1a cell lineage (19, 30). In adult mice, adipose CD5^+^ B cells appear to be segregated between the abdominal and the thoracic cavities. In addition to a self-renewal property described by earlier studies for B-1a cells (66), our studies show that the lifetime maintenance of CD5^+^ B cells in PATs also relies on a continuous supply through the circulation. Our finding that CD5^+^ B cells are enriched in PATs and reside in FALCs is reminiscent of prior studies on B-1 cells in FALCs of omentum and mesenterium (44). This is not particularly surprising, considering the shared developmental origin of these tissues from the splanchnopleuric mesoderm (67–69). Interestingly, it has been shown that this intraembryonic region contains progenitors for B-1a cells (70), and this observation has been extended to the fetal omentum (71). PATs may have evolved in the thoracic cavity as part of the surveillance mechanism for immunological and inflammatory insults, similar to omentum in the abdominal cavity (72). Additional investigation is needed to address whether IL-10-producing CD5^+^ B cells in PATs contain a distinct fraction of B-1a cells that can further expand upon pathological challenge. Alternatively, it remains possible that all B-1a cells in PATs are capable of proliferating and producing IL-10 upon exposure to particular signals. Similarly, it remains possible that subsets of B-2 cells such as CD5^+^ B10 cells contribute to the pool of IL-10-producing B cells in PATs. Regardless, our studies show that CD5^+^ B cells in PATs can function as a source of IL-10.

It has been reported that subsets of B cells that produce pro-inflammatory cytokines worsen the outcome of acute MI by promoting the early inflammatory phase and the recruitment of pro-inflammatory myeloid cells following MI (24, 25). Opposing effects of pro-inflammatory and IL-10-producing B cells have been observed for several chronic inflammatory diseases such as obesity (34, 73). Our studies now extend this concept to the pathophysiology of acute MI. Our findings have shown that IL-10-producing B cells regulate the recruitment of anti-inflammatory monocytes and the clearance of pro-inflammatory monocytes during the resolution phase post-MI. Additionally, it has been shown that IL-10 influences other downstream cellular targets of hematopoietic and non-hematopoietic origins to collectively improve the outcome of acute MI (15, 16). A better understanding of the interactions between IL-10-producing B cells and other cell types relevant to the progression of MI-induced inflammation may identify new targets for prevention and treatment of acute MI. Previous studies have revealed a lower efficiency for B cell progenitors to produce B-1a cells compared to B-2 cells during adulthood (49–51). Our findings suggest a similar scenario for the expansion of B cell subsets in PATs in response to acute MI, as reflected by a decrease in the percentage of CD5^+^ B cells among B cells. Considering the opposing roles for the different subsets of B cells in MI-induced inflammation, therapeutic strategies that promote IL-10-producing CD5^+^ B cells and/or suppress pro-inflammatory B cells may be worthwhile to explore.

Our studies indicate that IL-33 is preferentially expressed in PATs and contributes to the enrichment of and IL-10-producing capacity by CD5^+^ B cells under homeostatic conditions. Additional research is needed to examine the cellular and molecular mechanisms by which PAT-derived IL-33 leads to the expansion of CD5^+^ B cells. It is likely that a variety of cell types in PATs express IL-33, as was previously documented for IL-33 produced by abdominal VATs (52). The immediate cellular targets of IL-33 in PATs are also unclear. Innate lymphoid type 2 cells (ILC2s) are the best-studied IL-33-responding cells. In response to IL-33 signaling, these cells release type 2 cytokines such as IL-5 to influence downstream target cells (52). In addition to ILC2s, several other immune cell types can respond to IL-33 to release type 2 cytokines (52). This includes B-1 cells, in particular B-1b cells (54). As such, it remains possible that B-1b cells and/or ILC2s in PATs can respond to IL-33 to release type 2 cytokines. The latter might then act on CD5^+^ B cells, in a manner similar to the response of B-1a cells in PerC (74, 75).

The reported findings place IL-10-producing B cells in the anti-inflammatory response that terminates MI-induced inflammation to facilitate tissue repair. They also expand our understanding of the mechanisms underlying cardioprotection mediated by IL-33. Together, our findings identify IL-10-producing B cells as novel targets to improve the outcome of acute MI.

## Materials and Methods

### Mice

The following mouse strains of both sexes were used for this study. C57BL/6J (stock # 000664, WT B6, CD45.2^+^), B6 CD45.1 (stock # 002014, CD45.1^+^), IL-10 KO (stock # 002251) and μMT^-^ (stock # 002288) mice on the B6 background were from The Jackson Laboratory. IL-33 KO and ST2 KO mice on the B6 background were obtained from Dr. Michael Rosen at Cincinnati Children’s Hospital Medical Center with permission from Drs. Susumu Nakae (RIKEN Center for Life Science Technologies) and Andrew McKenzie (MRC Laboratory of Molecular Biology, Cambridge UK). Mice were fed with a regular chow diet (5LOD, Labdiet) and water ad lib, maintained on a 12-hour light/dark cycle under specific pathogen-free conditions, and examined at 10-15 weeks of age or at the ages specified for individual experiments. Age- and sex-matched mice between the experimental and control groups were used in each experiment.

### Reagents

We used fluorescently labeled antibodies purchased from BD Biosciences, Ebioscience, Biolegend, or Tonbo Biosciences. These include antibodies against mouse CD45, CD45.1, CD45.2, TCR-β, CD3, CD90, CD19, B220, CD5, CD43, CD11b, CD11c, F4/80, IgM, IgD, Ly-6C, Ly-6G, MHC class II I-A^b^, PanNK, NK1.1, IL-10, ST2, Siglec F and appropriate isotype controls for surface and intracellular labeling. Non-specific binding was blocked using anti-mouse CD16/CD32 (BD Biosciences or Tonbo Biosciences). Viability staining solutions 7-amino-actinomycin D (7AAD) and propidium iodide (PI) were from Ebioscience. We purchased phorbol 12-myristate 13-acetate (PMA), ionomycin, and *Escherichia coli* lipopolysaccharide (LPS) from Sigma-Aldrich. Collagenase type I was from Worthington Biochemical Corporation. Collagenase and DNAse for digesting LVs (see below) were obtained from Sigma-Aldrich. Cell culture medium and supplements, and PBS were from either Invitrogen Life Technologies or Corning Life Sciences. We used fetal bovine serum (FBS) from Atlanta Biologicals. Other reagents are described below in specific assays.

### Cell preparation

In experiments examining different adipose depots, or PATs and LVs, we harvested and analyzed tissues in parallel from each mouse. Tissues were rinsed in cold PBS, minced to paste-like consistency, and digested by enzyme solutions specified as follows. Adipose tissues were digested with collagenase I as described to yield SVFs (76). LVs were cleaned under a dissecting microscope, minced, and digested by an enzyme mixture containing collagenase I, collagenase XI, DNase I, and hyaluronidase dissolved in RPMI 1640 medium plus HEPES, as described (64). After digestion, the enzymes were removed by washing 2 times with RPMI 1640 medium containing 5% FBS. The digestions were then subjected to density centrifugation using Percoll (GE Healthcare Life Sciences) as described previously to purify leukocytes (76). PerC fluid was collected as described previously (34). Red blood cells were lysed using ACK buffer (Lonza).

### Surface and intracellular staining and flow cytometry

We performed surface and intracellular antibody staining as previously described (34, 76). Stained samples were analyzed on a BD FACSCanto II or BD LSR flow cytometer depending on the number of targeted markers. We characterized subsets of B cells and subsets of monocytes as described previously (34, 64). Intracellular staining of IL-10 was performed as described (34) on either freshly prepared cells or after *ex vivo* challenge (see below). The acquired data were analyzed using FlowJo software (Tree Star Inc.). In addition to the frequency, we calculated the total number of a cell subset per organ and the number of a cell subset per gram of tissue.

### Ex vivo culture of B cells

We performed *ex vivo* B cell functional assays to examine the capacity of B cell subsets to produce IL-10 as previously described (34). Briefly, purified SVFs from VATs or leukocytes from LVs were added into each well of a 96-well round-bottomed tissue culture plate. Cells were cultured for 5 hours in the presence of GolgiStop (BD Biosciences) at 37°C in a humidified cell culture incubator with 5% CO_2_. The un-challenged samples were cultured with GolgiStop alone, and the challenged samples were cultured with GolgiStop plus PIL (50 ng/ml PMA, 500 ng/ml ionomycin, and 10 μg/ml LPS). At the end of culture, cells were harvested for surface and intracellular labeling with antibodies as described above.

### Histology and immunostaining of PATs

At sacrifice, hearts were perfused with cold PBS. We harvested hearts and surrounding tissues and cleaned them in cold PBS under a dissecting microscope. PATs were then separated from hearts. Tissues were fixed in 10% formalin overnight at 4°C. For whole-mount staining, tissues were washed in PBS, flattened on a glass slide and stained with H&E. Stained tissues were cover-slipped and viewed under a Nikon AZ100M microscope. For paraffin-embedded sections, fixed tissues were flattened, dehydrated and embedded. Serial sections of 5μm thickness were used for H&E staining and immunostaining. For the latter, we followed a protocol of immunohistochemistry as described using the Vectastain ABC Kit (Vector Laboratories) (77). Antibodies were purchased from Biolegend or Ebioscience and are specified in Figures. For frozen sections, fixed tissues were washed in PBS, equilibrated in 30% sucrose dissolved in PBS overnight, flattened, and embedded in OCT medium (Sakura Finetek). Serial sections of 5µm thickness were used for H&E, Oil Red O, or immunofluorescence staining with antibodies specified in Figures. Immunofluorescence stained slides were cover-slipped using a mounting media containing DAPI (Vector Laboratories) to identify cell nuclei. All images were acquired using a Nikon AZ100M microscope, and viewed using NIH ImageJ software.

### Parabiosis

We performed parabiosis following a procedure previously described by other investigators (64). Age- and sex-matched congenic CD45.2^+^ and CD45.1^+^ mice were paired as parabionts. Surgery was performed using sterile techniques and a single joined pair was housed in a clean cage with food, including gel diet, and water readily accessible. Mice were sacrificed 4 weeks post-surgery. We analyzed both parabionts from each joined pair. Tissues were harvested to purify SVFs. B cell subsets were phenotyped by flow cytometry. The percentage of CD45.2^+^ or CD45.1^+^ cells among each B cell subset is presented for each parabiont.

### Adoptive transfer

To track transferred cells, we used donor cells from CD45.1^+^ mice. We collected B cells from PerC or purified SVFs from PATs, and injected cells into PerC of CD45.2^+^ mice. Recipients were sacrificed 3 or 7 days post-transfer. Donor-derived CD45.1^+^ cells were further analyzed for B cell subsets.

### B cell-specific deletion of IL-10

We employed a mixed bone marrow chimera approach to delete IL-10 specifically in B cells of B6 mice (59). Briefly, donor bone marrow cells were harvested from femurs and tibias of WT B6, IL-10 KO, or μMT^-^ mice at 8-12 weeks of age. Recipients were WT B6 mice at 6-8 weeks of age and were lethally irradiated 6 hours prior to transplantation. For generating B-IL-10 KO mice, recipients received intravenous injection of a mixture containing 20% of IL-10 KO and 80% of μMT^-^ bone marrow cells. For generating B-WT mice, recipients received intravenous injection of a mixture containing 20% of B6 WT and 80% of μMT^-^ bone marrow cells. Mice were allowed 8 weeks to reconstitute the transplanted bone marrow, and were then used for the subsequent experiments.

### Myocardial infarction

We induced acute MI in mice by permanently ligating LAD following a recently developed low-invasive procedure (57). Briefly, mice were anesthetized with isoflurane (2%). The chest was shaved, and a left thoracotomy was performed in the 4th intercostal space by making a small cut in the skin and bluntly opening the muscle wall. The heart was pushed out of the thoracic space to visualize the left ventricle. For myocardial infarction, the LAD was ligated at a site 3 mm from its origin using 8-0 sutures. For sham controls, no ligation was made. The heart was immediately placed back into the thoracic space followed by closure of the muscle wall. The skin was then closed with 6-0 sutures. Mice were allowed to recover from anesthesia and were monitored throughout the recovery period. They received subcutaneous injections of buprenorphine every 12 hours for 48 hours, starting 30 minutes before the surgery.

### Cardiac function and infarct size

Echocardiography under anesthesia was employed to assess the function of LVs as described (63), using Vevo 2100 (VisualSonic, Toronto, Canada). Briefly, mice were anesthetized with isoflurane inhalation (1.5%). Two-dimensional echocardiographic views of the mid-ventricular short axis were obtained at the level of the papillary muscle tips below the mitral valve. LV wall thickness and internal dimensions were measured. EF and FS were calculated from the parameters taken from M-mode images. Masson’s trichrome stain of collagen was used to visualize MI scar. At sacrifice, hearts were harvested after perfusion with cold PBS, cleaned under a dissecting microscope, and fixed in 10% formalin overnight at 4^0^C with rocking. Hearts were then cut into halves along the long axis to display the 4 chambers and embedded in paraffin. Sections of 8μm thickness were used for Masson’s trichrome stain using a kit purchased from Sigma-Aldrich. Images were taken using a Nikon AZ100M microscope. MI scar was measured using NIH ImageJ software. Scar size was expressed as percentage of LV circumference.

### Real-time PCR

At sacrifice, harvested tissues were snap-frozen in liquid nitrogen and stored at −80°C. Total RNA was extracted using Trizol reagent (Invitrogen Life Sciences), then reverse transcribed into cDNA using Superscript III First-Strand Synthesis kit (Invitrogen Life Sciences). Real-time PCR was performed as previously described using SYBR green and a Bio-Rad CFX thermal cycler (34, 76). β-actin was used as the internal control. The primer sequences are: TNF-α forward, 5’-ACGGCATGGATCTCAAAGAC-3’; TNF-α reverse, 5’-AGATAGCAAATCGGCTGACG-3’; MCP-1 forward, 5’-CCACTCACCTGCTGCTACTCA-3’; MCP-1 reverse, 5’-TGGTGATCCTCTTGTAGCTCTCC-3’; β-actin forward, 5’-TACAGCTTCACCACCACAGC-3’; β-actin reverse, 5’-AAGGAAGGCTGGAAAAGAGC-3’; IL-33 forward, 5’-ATGGGAAGAAGCTGATGGTG-3’; IL-33 reverse, 5’-CCGAGGACTTTTTGTGAAGG-3’; ST2L forward, 5’-AAGGCACACCATAAGGCTGA-3’; ST2L reverse, 5’-TCGTAGAGCTTGCCATCGTT-3’; sST2 forward, 5’-TCGAAATGAAAGTTCCAGCA-3’; sST2 reverse, 5’-TGTGTGAGGGACACTCCTTAC-3’; IL-18 forward, 5’-AGTGAACCCCAGACCAGACTGA-3’; IL-18 reverse, 5’-CCCTCCCCACCTAACTTTGATGTA-3’.

### Statistics

Data are presented as means ± SEM. Statistical analyses were performed using unpaired 2-tailed t test or One-way ANOVA with post-hoc Tukey HSD test after examining distribution normality of the data. A *p* value less than 0.05 was considered statistically significant.

### Study approval

All procedures for use of mice in the study were approved by the Institutional Animal Care and Use Committee at Vanderbilt University School of Medicine.

## Acknowledgements

This work was supported by NIH grants R01DK081536 (L.W. and L.V.K.), R01DK104817 (L.V.K.), R01HL133290 and R01HL119234 (H.L.). R.D. and C.C. were medical students from New York Institute of Technology College of Osteopathic Medicine and Rutgers Robert Wood Johnson Medical School, respectively, who participated in the NIH/NIDDK-sponsored Vanderbilt Medical Student Research Training Program (T35DK07383). J.L.P. was supported by institutional training grants (T32HL069765 and T32AR059039). FACS sorting was performed in the Flow Cytometry Shared Resource at Vanderbilt University Medical Center, supported by the Vanderbilt Ingram Cancer Center (P30 CA68485) and the Vanderbilt Digestive Disease Research Center (DK058404). Imaging was performed in the Cell Imaging Shared Resource at Vanderbilt University Medical Center, supported by the Vanderbilt Diabetes Research and Training Center (DK020593) and the Vanderbilt Ingram Cancer Center (P30 CA68485). We thank Drs. Michael Rosen, Susumu Nakae, and Andrew McKenzie for providing IL-33 KO and ST2 KO mice on the B6 background.

Author contributions
L.W. designed research; L.W., R.D., C.C., J. L.P., Q. Z., and Z.W. performed research and analyzed data; H.L. participated in the design and data analysis for cardiac experiments; L.W. and L.V.K. wrote the paper.

