## supplemental table and figures for "IL-10-producing B cells are enriched in murine pericardial adipose tissues and ameliorate the outcome of acute myocardial infarction"

#### Supplemental Figures

**Supplemental Figure 1. Flow cytometry gating of B cell subsets.** Cells were harvested from the indicated locations of adult WT B6 mice and analyzed by flow cytometry. **a**. Gated B cells were separated into B-1 and B-2 cells and further divided into B-1a and B-1b cells based on cell surface markers. **b**. $CD5^+$ B cells were gated among B cells based on surface CD5 expression.

**Supplemental Figure 2. Detection of IL-10 in $CD5^+$ B cells. a**. SVFs were harvested from PATs of adult WT B6 mice. Intracellular IL-10 was examined in freshly isolated $CD5^+$ B cells by flow cytometry (FMO: fluorescence minus one). **b**. SVFs from the indicated VAT depots of mice with the indicated genotypes were cultured for 5 hours in the presence of monensin, plus or minus the mitogen mixture PIL (PMA, ionomycin, and LPS). Cells were then examined for the intensity of IL-10 staining in $CD5^+$ B cells. Summary of 2 independent experiments is shown (n=6). $^{\#}$: p<0.05; and *: p<0.01.

**Supplemental Figure 3. Surface phenotype of $CD5^+$ B cells.** Cells were harvested from adult WT B6 mice. Subsets of B cells in SVFs of PATs were labeled with the indicated surface markers and analyzed by flow cytometry. Representative flow cytometry plots are shown (n=4 from 2 independent experiments).

**Supplemental Figure 4. Population dynamics of $CD5^+$ B cells in PerC and periaortic VATs.** Prevalence of $CD5^+$ B cells in the indicated locations was examined in

WT B6 mice at different ages. A summary of 2 independent experiments (n=8 for each age group) is shown.

**Supplemental Figure 5. Histology and immunostaining of PATs and FALCs**. Tissues were harvested from adult WT B6 mice. **a**. Whole-mount PATs together with pericardium were stained with H&E and photographed (magnification: 0.5x). Two areas with densely packed cells are enlarged to show clusters (magnification: 4x). **b**. Paraffin-embedded sections of PATs together with pericardium were stained with H&E and photographed (magnification: 0.5x). Four areas are enlarged to show different types of adipocytes (magnification: 10x). **c**. Frozen sections of PATs together with pericardium were stained with either H&E or immunostained for the indicated markers and photographed (magnification: 0.5x). One area containing a FALC is enlarged to illustrate different lymphocytes (magnification: 20x).

**Supplemental Figure 6. Histology and immunostaining of PATs and FALCs in IL-10 KO mice**. Tissues were harvested from mice of the indicated genotypes. Serial paraffin-embedded sections of PATs together with pericardium were stained with either H&E (magnification: 0.5x; upper panels) or for a B cell marker (lower panels), and were then photographed. Two areas are enlarged to show B cells in FALCs (magnification: 20x).

**Supplemental Figure 7. Maintenance of $CD5^+$ B cells in perigonadal VATs during adulthood.** Congenic $CD45.1^+$ and $CD45.2^+$ mice were joined by parabiosis and examined 1 month later. Subsets of B cells in perigonadal VATs of each parabiont were examined by flow cytometry. Summary of 3 independent experiments with 11 pairs of parabionts is shown. *: p<0.01; and ns: not statistically different.

**Supplemental Figure 8. Influence of IL-33 signaling on subsets of B cells in PATs**. Adult WT B6 and IL-33 KO mice (**a**), or WT B6 and ST2 KO mice (**b**) were sacrificed to prepare SVFs from PATs. Cells were then examined by flow cytometry for the prevalence of B cell subsets. Summary of 2 independent experiments is shown (n=9-12 for each group). *: p<0.01.

**Supplemental Figure 9. MI-induced response in PATs**. WT B6 mice underwent MI or sham operation and were analyzed 5 days post-surgery. Whole-mount PATs together with pericardium were stained with H&E and photographed (magnification: 0.5x). Two areas in each treatment with densely packed cells are enlarged to show clusters (magnification: 4x).

**Supplemental Figure 10. Flow cytometry gating of B cell subsets in LVs.** Cells were harvested from the indicated tissues of adult WT B6 mice and analyzed by flow cytometry. **a**. Gated B cells were separated into B-1 and B-2 cells and further divided into B-1a and B-1b cells based on cell surface markers. **b**. $CD5^+$ B cells were gated among B cells based on surface CD5 expression.

**Supplemental Figure 11. IL-10 production in B cells freshly purified from PATs and LVs**. WT B6 mice underwent MI or sham surgery. Mice were examined 8 days post-surgery. Intracellular IL-10 was examined in B cells freshly purified from PATs or LVs by flow cytometry. Representative flow cytometry plots from 2 independent experiments are shown (n=4-6 per group).

**Supplemental Figure 12. Detection of IL-10 in B cells of B-WT and B-IL-10 KO mice**. SVFs were harvested from PATs of mice with the indicated genotypes. Cells were cultured for 5 hours in the presence of monensin, plus or minus the mitogen mixture PIL (PMA, ionomycin, and LPS). $IL\text{-}10^{hi}$ B cells were examined by flow cytometry. Representative flow cytometry plots from 2 independent experiments are shown (n=6).

**Supplemental Figure 13. Murine echocardiography**. Mice underwent MI surgery and were examined for LV function by echocardiography 3 weeks after surgery. A representative M-mode image with measurements is shown.

**Supplemental Figure 14. Flow cytometry analysis of monocyte subsets among leukocytes of LVs**. B-IL-10 KO and B-WT mice underwent MI surgery. Mice were sacrificed at the indicated time point post-MI to purify leukocytes from LVs. Subsets of monocytes were analyzed by flow cytometry. Representative flow cytometry plots are shown.

**Supplemental Table 1: B cell subsets in adipose depots and in left ventricles**

| | B cells | B-2 cells | B-1 cells | B-1a cells | B-1b cells |
|---|---|---|---|---|---|
| | (% among SVFs) | (% among B cells) | (% among B cells) | (% among B-1 cells) | (% among B-1 cells) |
| BAT | 3.94 ± 0.51 | 90.66 ± 1.16 | 8.24 ± 1.18 | 23.35 ± 2.45 | 75.85 ± 2.42 |
| SAT | 1.54 ± 0.21 | 88.76 ± 2.35 | 9.72 ± 2.26 | 32.44 ± 3.59 | 66.68 ± 3.66 |
| Perigonadal | 1.39 ± 0.27 | 61.3 ± 2.19 | 37.53 ± 2.21 * | 45.28 ± 4.73 | 54.25 ± 4.77 |
| Retroperitoneal | 1.19 ± 0.35 | 60.99 ± 3.88 | 37.66 ± 3.86 * | 53.96 ± 1.5 | 45.68 ± 1.52 |
| Mesenteric | 1.58 ± 0.34 | 65.24 ± 7.56 | 32.9 ± 7.7 * | 49.83 ± 3.59 | 49.73 ± 3.55 |
| Periaortic | 12.44 ± 1.44 | 59.74 ± 3.59 | 39.03 ± 3.38 * | 53.9 ± 4.66 | 45.53 ± 5.52 |
| PATs | 26.15 ± 2.23 | 52.81 ± 6.24 | 42.13 ± 5.63 * | 57.68 ± 1.11 | 42.03 ± 1.11 |
| Ventricles | 6.17 ± 0.99 ** | 92.05 ± 0.64 | 6.65 ± 0.56 ** | 9.42 ± 0.92 ** | 87.53 ± 1.19 |

Data are presented as mean ± SEM of 8 mice in 2 experiments.

*: Significantly higher compared to those in BAT and SAT ($p<0.01$)

**: Significantly lower compared to those in PATs ($p<0.01$)

### Supplemental Figure 1

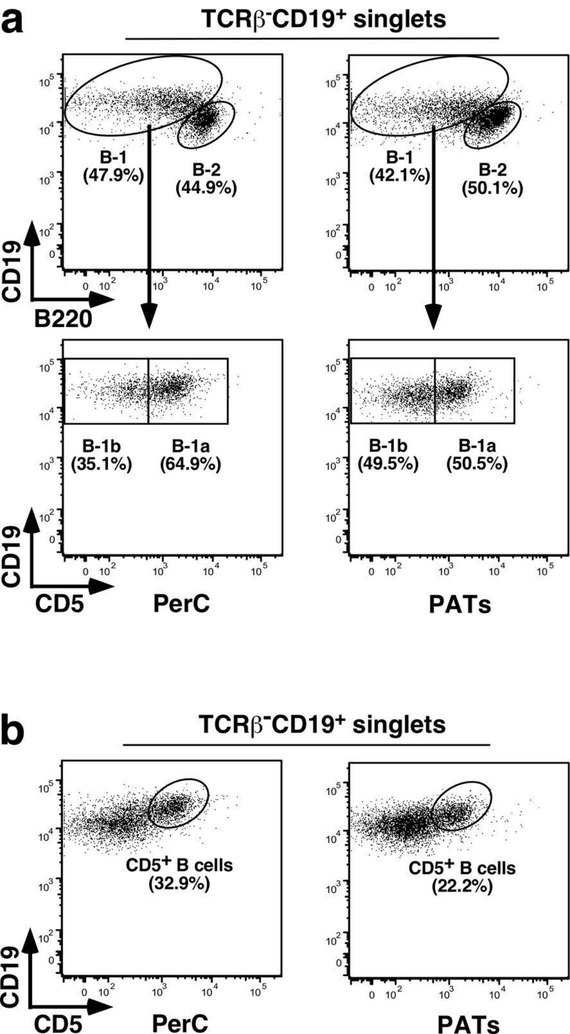

### Supplemental Figure 2

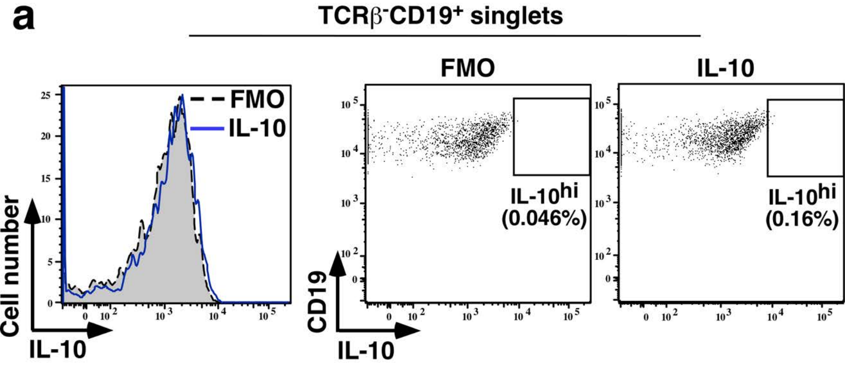


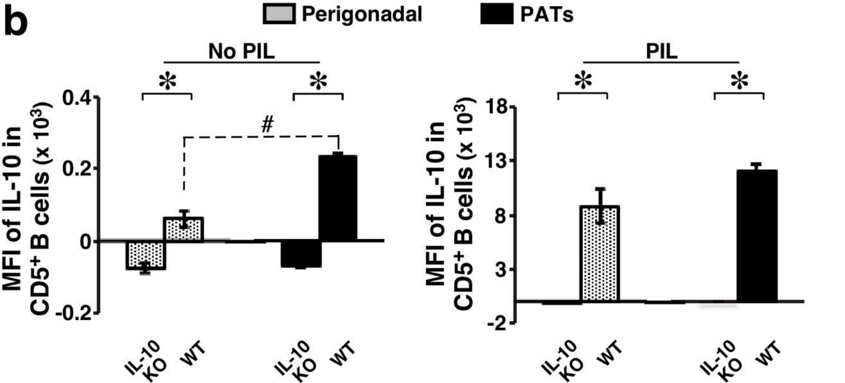

### Supplemental Figure 3

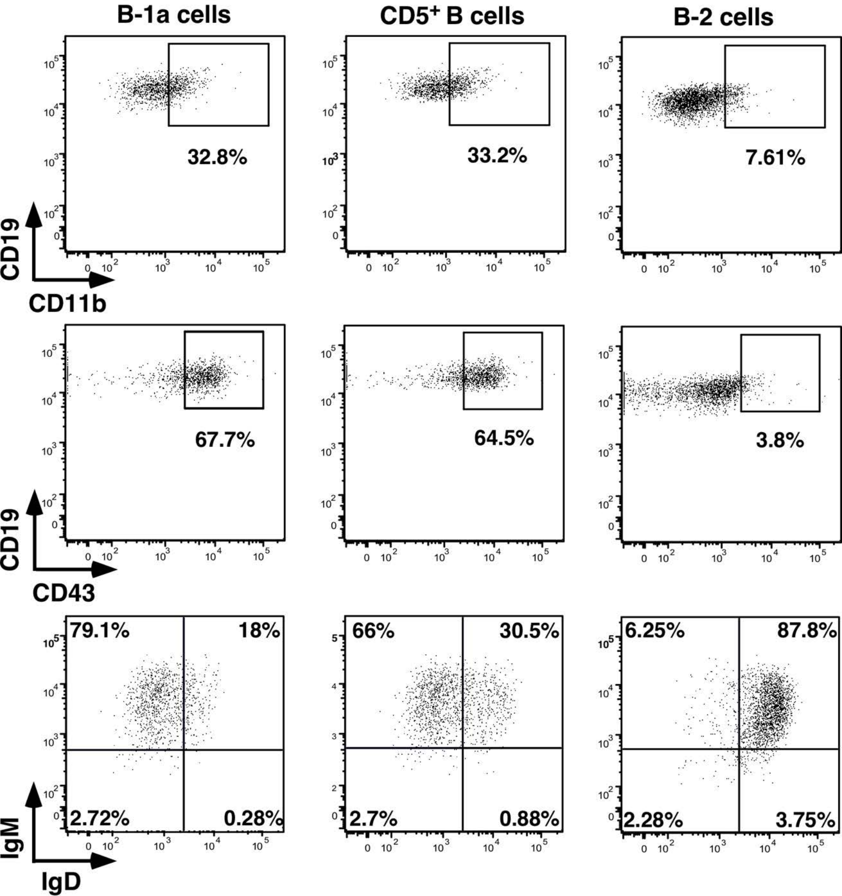

### Supplemental Figure 4

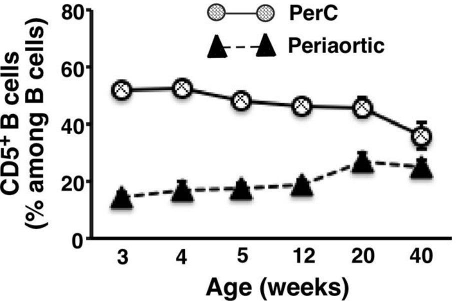

### Supplemental Figure 5

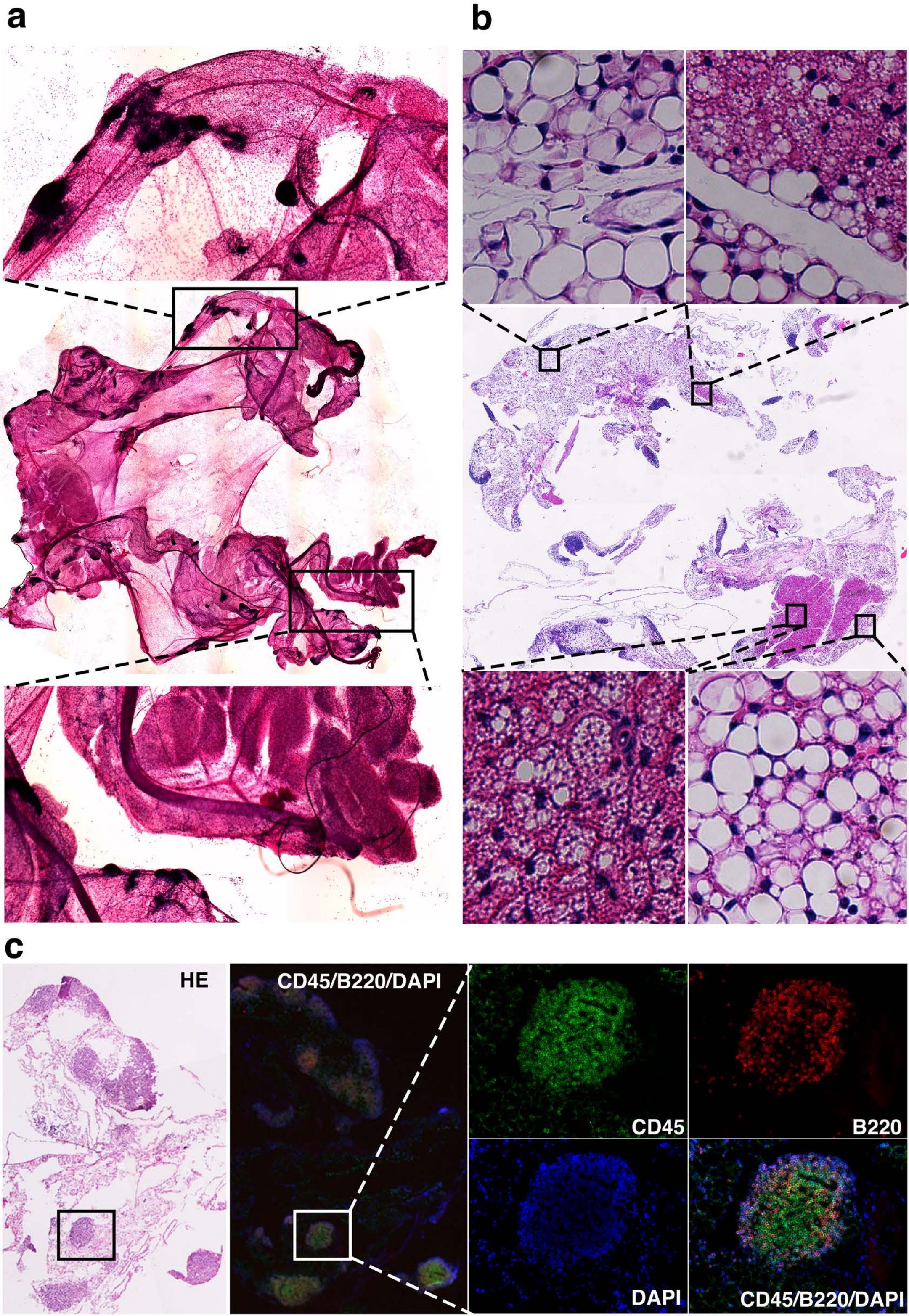

### Supplemental Figure 6

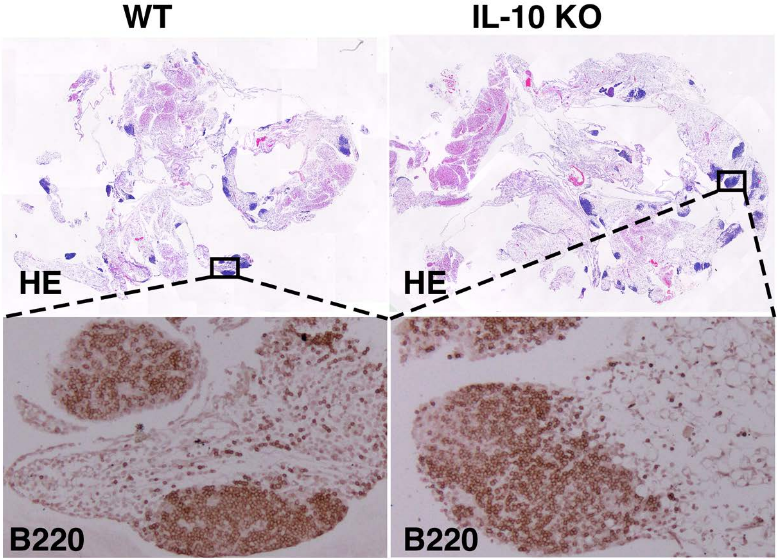

### Supplemental Figure 7

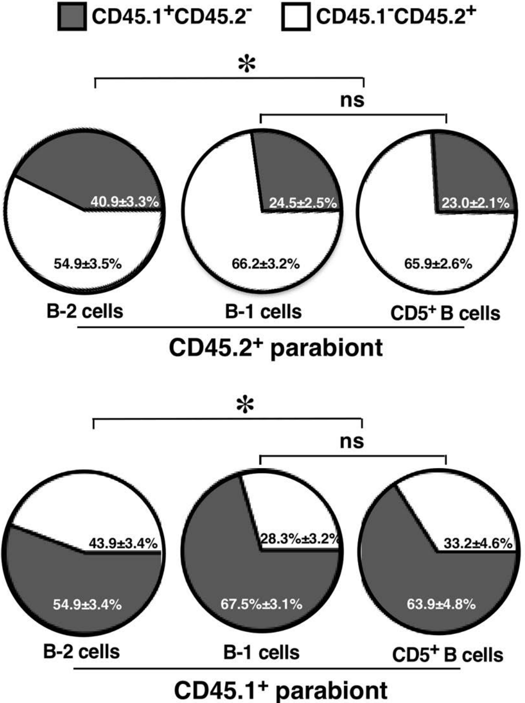

### Supplemental Figure 8

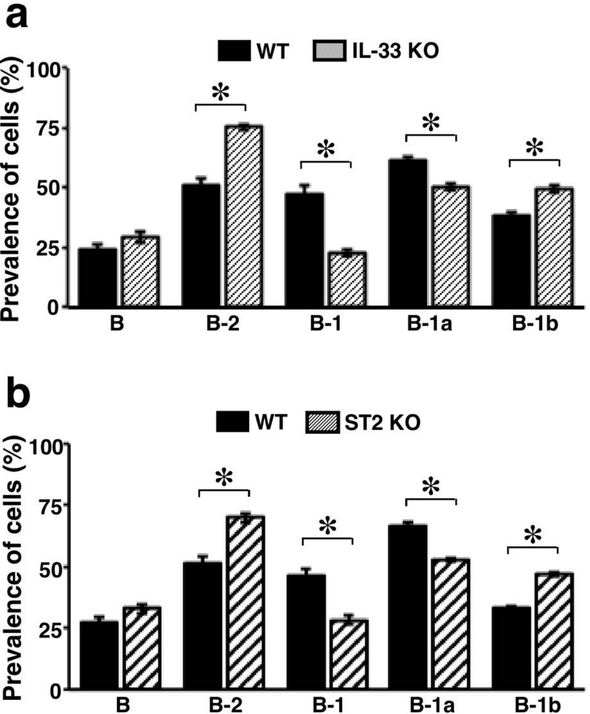

### Supplemental Figure 9

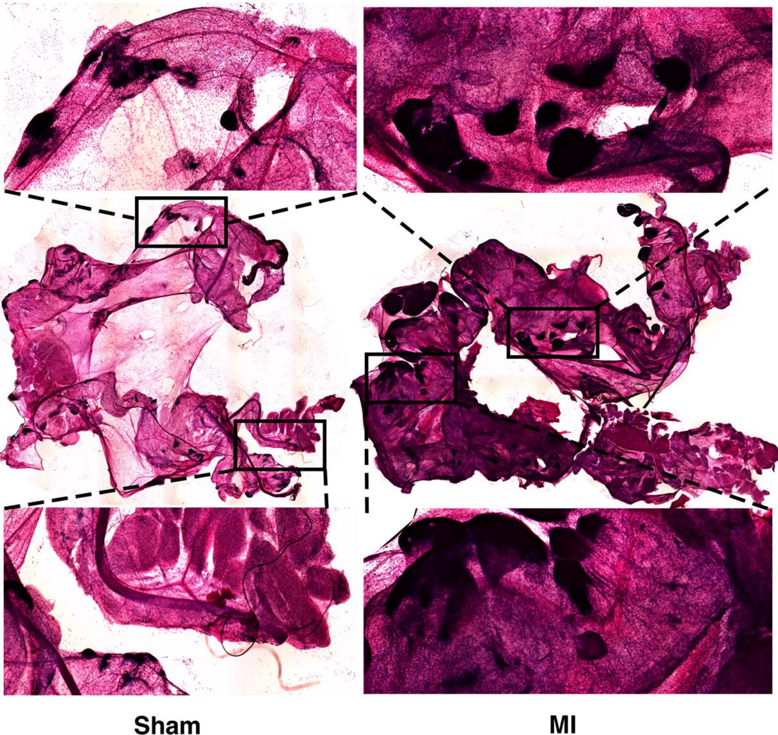

### Supplemental Figure 10

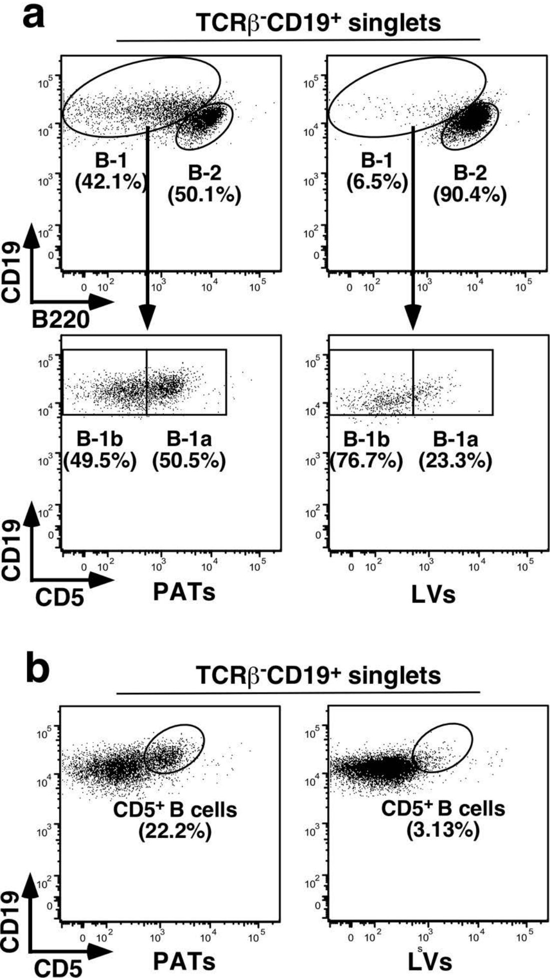

### Supplemental Figure 11

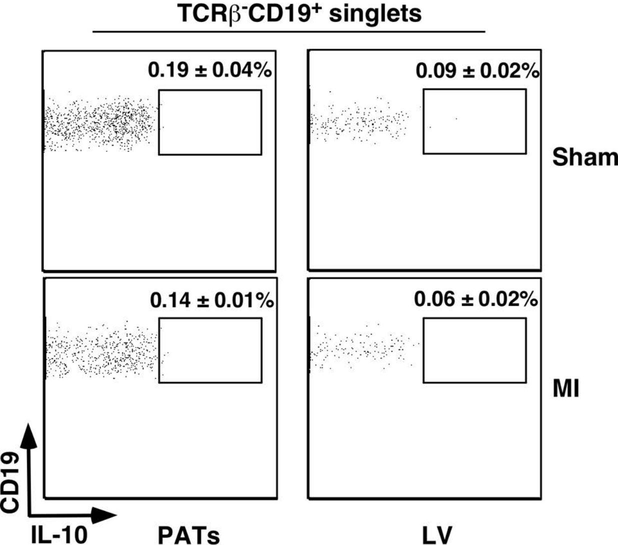

### Supplemental Figure 12

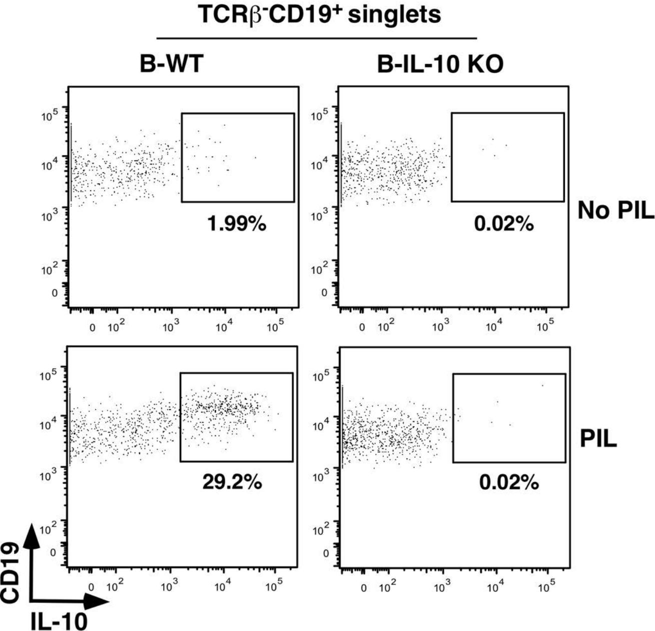

### Supplemental Figure 13

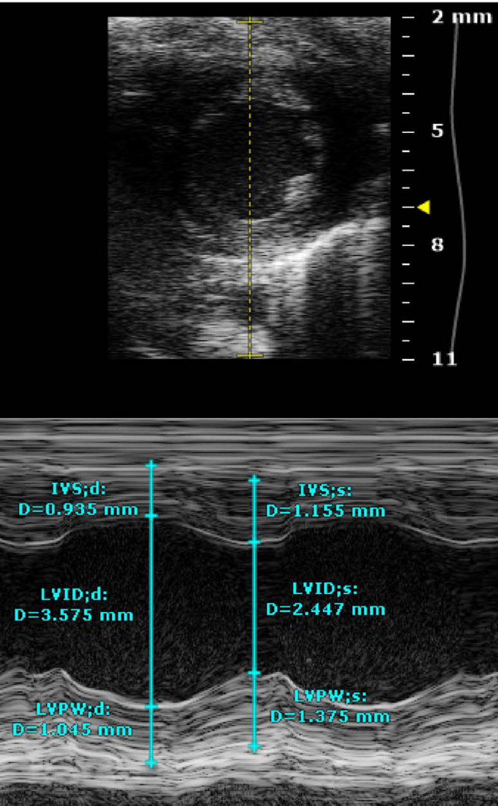

### Supplemental Figure 14

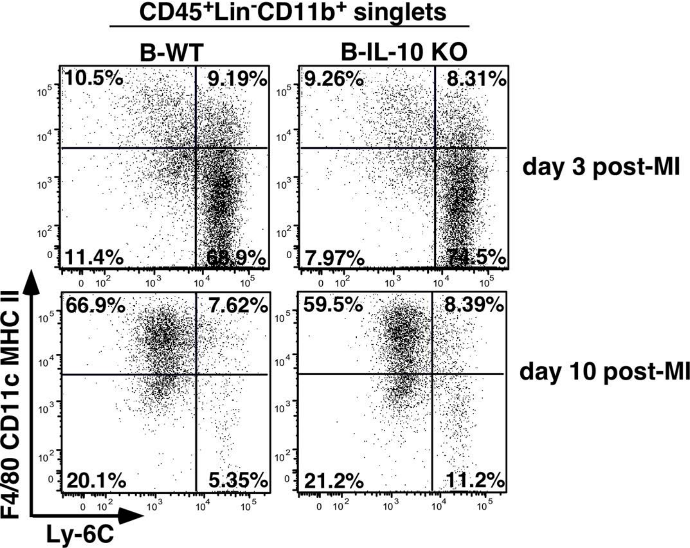
